# Assessing the predation function via quantitative and qualitative interaction components

**DOI:** 10.1101/2020.05.16.099721

**Authors:** Carlos Martínez-Núñez, Pedro J. Rey

## Abstract

Interactions among organisms can be defined by two main features: a quantitative component (i.e. frequency of occurrence) and a qualitative component (i.e. success of the interaction).

Measuring properly these two components at the community level, can provide a good estimate of the ecosystem functions mediated by biotic interactions. Although this approach has been frequently applied to evaluate the eco-evolutionary consequences of mutualistic relationships, it has never been extended to the predation function and the associated pest control ecosystem service.

Here, we introduce a simple measure that accounts for the quantitative and the qualitative components of predation interactions, and facilitates a precise characterization of this ecosystem function at the community level, while accounting for variations at species and individual levels.

This measure arises as a fine indicator of predation pressure, and provides great opportunities to better understand how different components of predation and pest control potential vary across environmental gradients.

## Introduction

The assessment of ecosystem functions and their relation to abundance and biodiversity is a central topic in ecology (Hooper et al., 2005; Woodcock et al., 2019). Under the current scenario of Global Change, predicting variations in these ecosystem functions is a key challenge for ecologists and human societies (Ruckelshaus et al., 2020).

Ecosystem functions have traditionally been estimated through the abundance or diversity of the individuals/species/traits potentially responsible for the specific function under study. Although this approach enables predictions, it has a number of shortcomings, because: i) the function provided by different species is usually not measured; ii) it assumes a similar contribution to the function by all the species belonging to the same group (e.g. pollinators, functional groups), or iii) at best, methods for predicting variations in the functions using central or dispersion (divergence) measures of functional traits in the community are used, masking the contribution of each species; and iv) conspecific individuals are deemed to have the same impact on the function despite some studies have shown that phenotypical and behavioral variability among genetically similar individuals can be important (Bruijning, Metcalf, Jongejans, & Ayroles, 2020).

Research on plant-animal mutualisms solved part of these problems by introducing two concepts grounded on an interaction approach, to assess the mutualistic function: the *quantitative component* (i.e. frequency of the interaction), and the *qualitative component* (i.e. interaction success for delivering the function). This approach has been more recently summarized (Schupp et al., 2010; 2017) as the *mutualism effectiveness framework*, also focused on other mutualistic interactions. According to this framework, variations in any of the two components could lead to changes in the ecosystem function under study (e.g. pollination or seed dispersal). This method, beyond species abundance, allows to include the interspecific variability in the effectiveness for delivering a function (i.e. changes in the quality component of the interaction) (e.g. Herrera, 1987), and also accounts for species-specific and individual-level information.

The partition of the function into quantity and quality components has been applied to address the effectiveness of pollinator assemblages (Rodríguez-Rodríguez, Jordano, & Valido, 2013), the environmental impacts on avian seed dispersal (Rey & Alcántara, 2014), and even the variation on pollination and seed-dispersal effectiveness across perturbation gradients simultaneously (Fontúrbel, Jordano, & Medel, 2017). Moreover, the two-dimensional representation of all possible combinations of the quantitative and qualitative components in a community, known as “*effectiveness landscapes*” (Schupp et al., 2010), has proven useful. Some authors have used it to study the functional equivalence or contribution of particular groups (González-Castro, Calviño-Cancela, & Nogales, 2015) or species (Blendinger, 2017) to different functions. For instance, Hervías-Parejo & Traveset (2018) compared the pollination effectiveness of insects and opportunistic Galapagos birds. Although this framework has been very fertile and useful in studying mutualistic functions, to the best of our knowledge, these concepts and methods have surprisingly not permeated to non-mutualistic interactions yet.

Here, we adopt the concepts developed in the mutualism effectiveness framework (Schupp et al., 2017) and extend them to antagonistic interactions. Specifically, we propose a method to assess the predation function by accounting for the qualitative component of interactions, at the species level, including intraspecific variations (individual level). We extrapolate it then to the whole community, using a quantitative component, to achieve a highly resolved measure of the community-wide predation function. Importantly, the measure we present in this work has wide applications to advance the study of the pest control ecosystem service. Last, we showcase these ideas and apply the introduced method to a case study, in which we compare the ant-mediated predation function against a major olive tree pest across landscape complexity anthropogenic gradients.

## Materials and Method

### Quantifying and decomposing the predation function

We propose a simple equation (Eq. 1) to calculate and predict the total predation ecosystem function provided by a species or community by extending the mutualism effectiveness framework (Schupp et al., 2017). In this case, the predation function is also measured by using two components: a quantitative and a qualitative component (see corresponding sections for a deeper description). Interestingly, independent variations in any of these two components imply changes in the predation function (see Box 1).

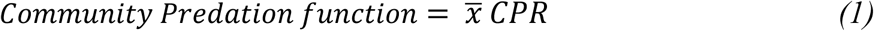

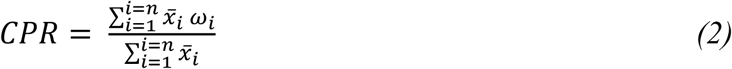

The quantitative component (first term in the equation) represents the abundance of the predator species guild being considered (i.e., the potential predators). It is here represented by the mean abundance per sample 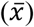 of the potential predators (although it could also be other central measure or the total sum of predator individuals in the community), which multiplies the Community Predation Response (CPR), the qualitative component. CPR is the expected probability of predation after encounter with a prey by a randomly chosen predator individual belonging to the predator guild (i.e. the studied group or species). The qualitative component is represented by the second term in the equation. Mathematically (Eq. 2), equals the mean abundance of each predator species (from 1 to *n* potential predator species) multiplied by its species’ specific response to prey encounter (*ω*_i_)(e.g. proportion of positive responses after encounter) weighted by the relative abundance of each species 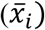 in the community 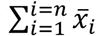. Likewise, the predation effectiveness of each species against a certain prey might be quantified with: 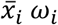 (abundance multiplied by its species-specific response).

### Quantitative component: Probability of encounter

The quantitative component represents the abundance of potential predators (or a guild of predators) in a site, which is closely linked to the probability of encounter and predation pressure (provided constant appetence). For most systems, especially those typically distant from ecological equilibrium, such as most agroecosystems, it is an easy assumption that probability of encounter and predation, increases linearly with abundance (Eq. 1). However, more complex relationships between the two components can be easily derived (e.g. logistic or exponential) if the specific system assumptions suggest it (e.g. to accommodate different Holling’s predator functional responses). Because density dependent processes such as territoriality, competence or closeness to the system carrying capacity (K) are expected to affect abundance, the Eq. 1 does account for them indirectly, through this quantitative component.

### Qualitative component: weighted species-specific response

The qualitative component measures how optimum the community assemblage or the interaction is to perform a certain function. In our framework, it represents the expected appetence for the prey, or predation success. It is important to note that this factor accounts for variations at the species and individual level. The specific response of different individuals is implicit at the species or group level (e.g. average or median of the response) as the probability of predation success after encounter (summarized as proportion of successful/unsuccessful encounters, for instance). The specific response of a species can be assessed experimentally as the proportion of positive responses to offered preys (e.g., Garrido et al., 2002, for a similar approach using diaspore offerings for ant-mediated seed dispersal). It also accounts for the predator species composition and relative abundance in the community which, independently of the quantitative component, may induce changes among sites in the predation function and pest control service. Hence, the qualitative component represents the expected predation probability by a randomly chosen potential predator in a community, after predator-prey encounter.

In the case that predation function largely depends on the predator’s probability to actively find the prey and/or the experimental quantification of positive/negative responses is impractical (e.g. offerings are not possible and only positive responses, or predation events, are detected), the *ω*_i_ parameter becomes the frequency of positive responses by species *i* (i.e. successful predation events). Because this relationship is not based on proportions (i.e. positive interactions from total), the measure should be corrected accounting for prey availability.

Also, if specific responses are known to be always positive and successful (e.g. 100% appetence for the prey), the predation function would depend mainly on the probability of encounter. Therefore, modelling variations of the quantitative component might be a good proxy.

## Results

The methods and concepts here introduced are showcased by means of a case study, where we compared the pest control ecosystem service provided by the ant community against a major olive tree pest, the olive moth (*Prays oleae*, Bernard), across gradients of landscape heterogeneity.

### Landscape effects on the ant-mediated pest control against the olive moth

Martínez-Núñez et al., 2020 (under review), sampled during seven months (from April to November), the ant community abundance at the farm scale (the quantitative component of the interaction) in 40 olive groves spanning a wide gradient of human-shaped landscape complexity at the regional scale. The authors also assessed ant species specific responses against the olive moth by offering real pest larvae to 25 foraging ant species, and calculated the qualitative component of the predation function at the community level (Table S1 in Appendix 2). A positive response was recorded when the ant interacted with the pest (i.e. touching it with the antennae) and attacked it. In contrast, the ant ignoring the larvae after contacting it was deemed as a negative response. Accounting for species abundance and the specific response of each species, the ant community predation function (i.e. expected predation pressure by ants over the pest) was calculated for each farm, using the Eq. 1. Here, we relate farm variation in community predation function to six metrics of compositional and configurational heterogeneity. The landscape metrics were obtained from a buffer of 1km radius around the centroid of each farm are: largest patch size (LPS), edge density (ED), distance to nearest neighbor (NDD), proportion of semi-natural habitat (SNH), patch richness (PR) and patch diversity (PD). For methodological details, see Appendix 1 in Supporting Information. We also represent the position of each species in the *predation function effectiveness landscape* using the “*effect.lndscp”* package (Jordano, 2014) in R (R Core Team, 2019), to assess the specific contribution of each species to the overall ant pest control service 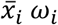. Last, we explore the existence of independent gradients in the variation of each component across landscape gradients, which can help to pinpoint the processes involved in the loss of the predation function.

Results show that the ant predation function over the olive moth larvae is higher in landscapes with a higher PR, higher PD, higher ED, intermediate LPS and NND (see Table 1). The Fig. 1, represents the position in the effectiveness landscape plot of each olive farm, according to their two components. This figure suggests that ant community responses (appetence for the prey) against the olive moth are generally high (i.e., in most farms there is a positive probability that a randomly chosen ant predates on the olive moth larvae), and there is a greater margin of community predation function improvement by increasing ant abundance. Interestingly, the quantity and quality components at the community scale, showed an overall similar response to landscape complexity (Fig. 2), although effects were more manifest for the quantitative component. This response was weak but negatively affected (i.e. less predation function) by landscape simplification. Both results could be expected from scenario *a* in Box 1 for which land use intensification and landscape homogenization would drive a decrease in predator abundance rather than in their response (qualitative component).

**Table 1:**
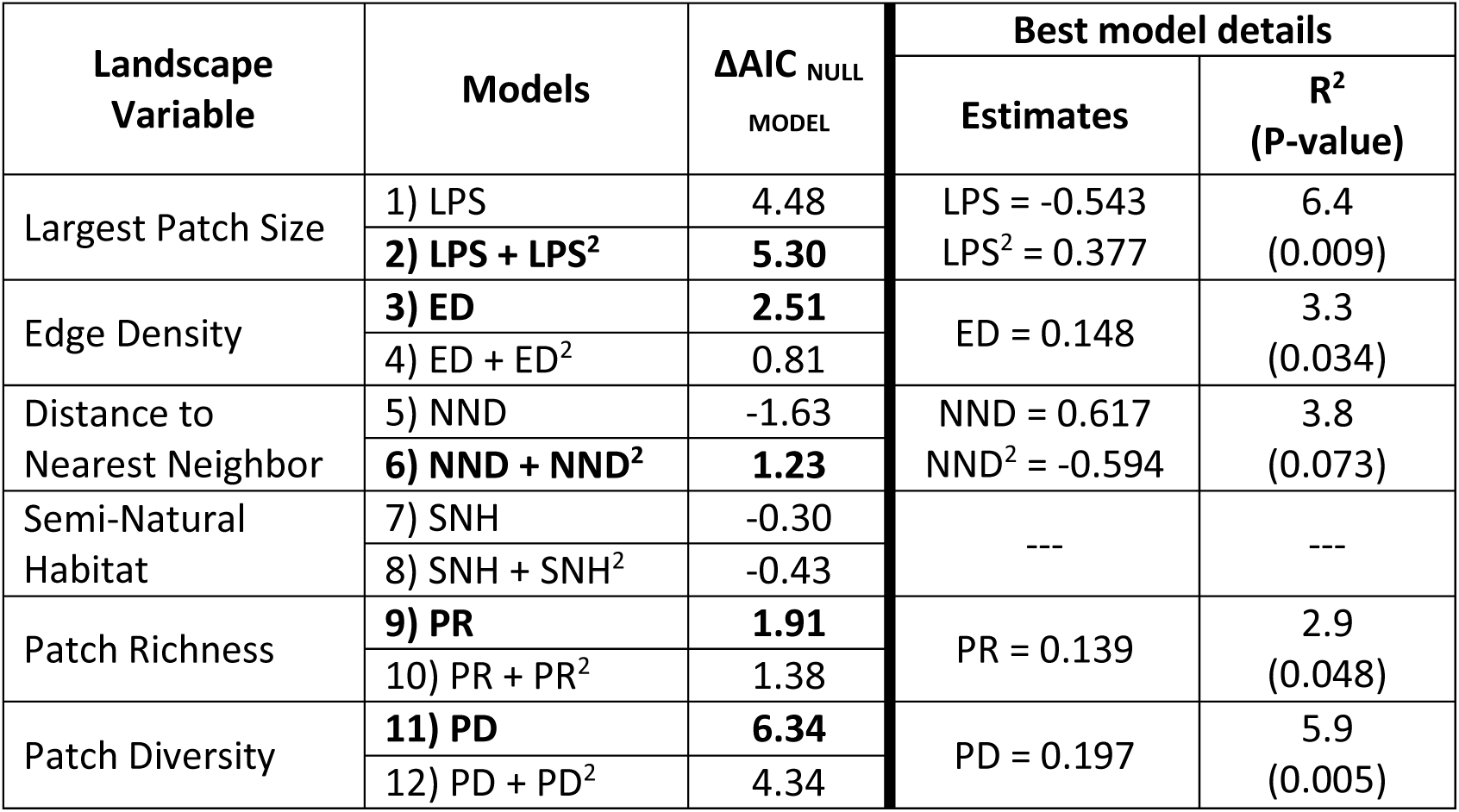
Response of Ant Community Predator Function (APF) to landscape variables: LPS (Largest Patch Size), ED (Edge Density), NND (Nearest Neighbor Distance), SNH (Semi-natural Habitat), PR (Patch Richness) and PD (Patch Diversity). Results from linear and quadratic terms, using linear mixed models, with olive farm as random factor. The best model (according to AIC, compared to the null model) is highlighted in bold. Terms’ estimates, marginal R^2^ and model P-value are shown for the best model (if any).

**Figure 1:**
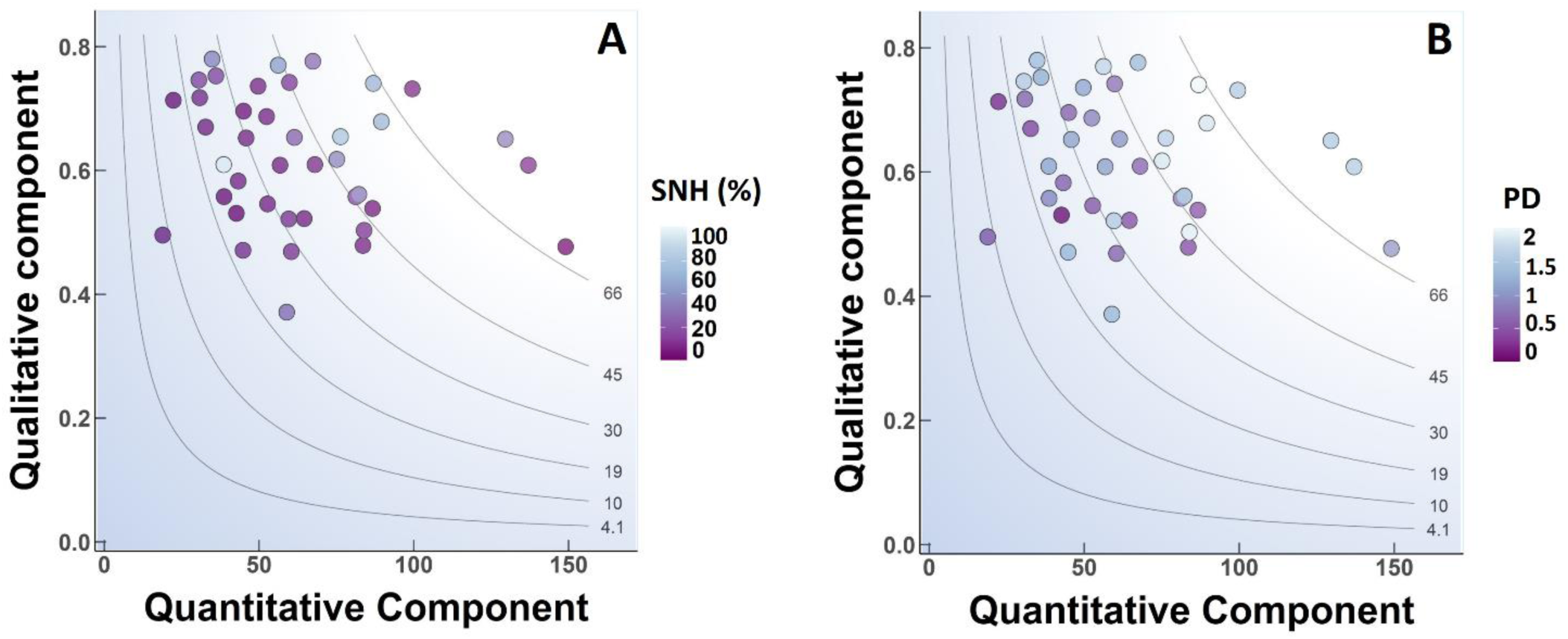
The predation effectiveness landscape in different sites (i.e. farms). Here, the quantitative component refers to mean abundance per pitfall trap. Isoclines represent all possible combinations of the quantitative and qualitative components that render the same total predation effectiveness. Point colors symbolize the amount of semi-natural habitat (SNH) in A, and Patch Diversity (PD) in B, surrounding the farms (1km radius scale).

**Figure 2:**
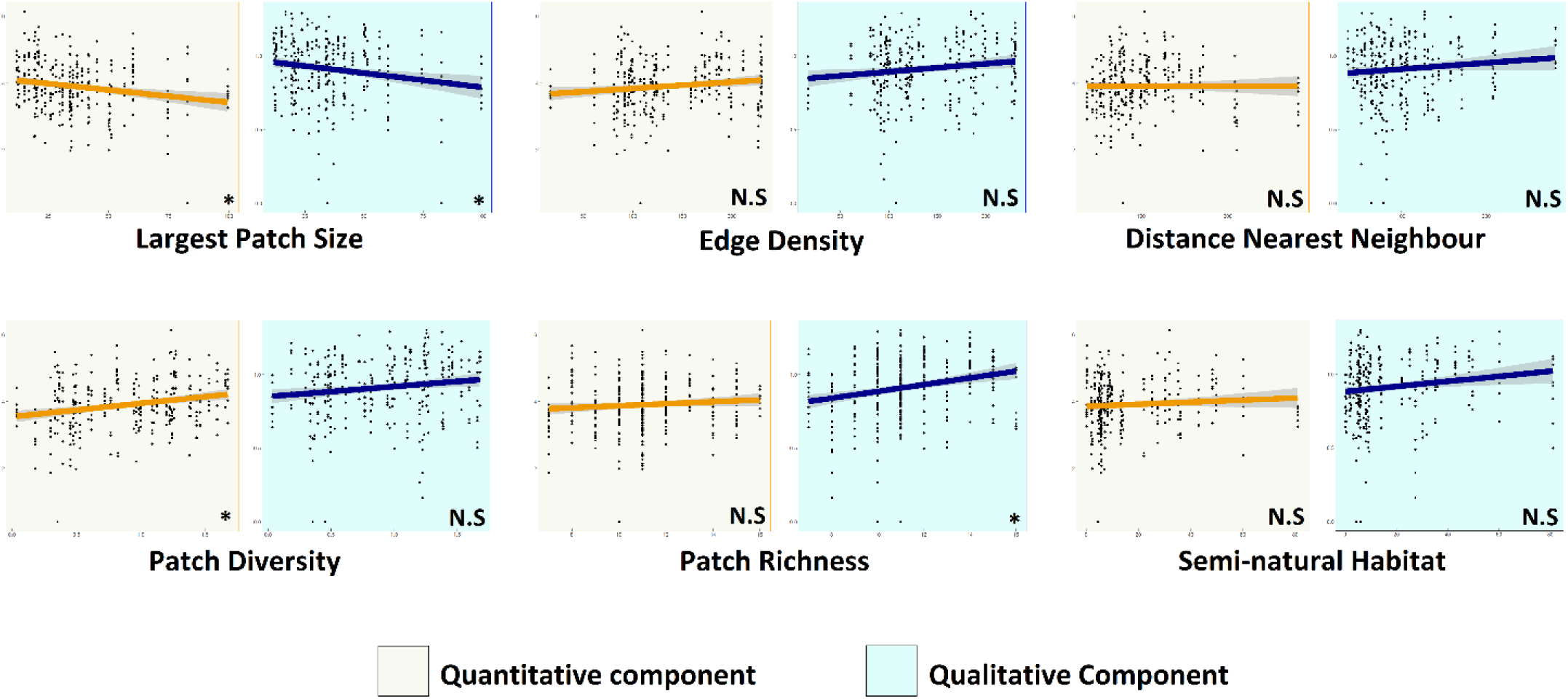
Variations of the quantitative and qualitative components across landscape complexity gradients. Linear mixed models, with landscape metric as fixed effect variable, and farm ID as a random effect variable. N.S = Non-significant model. * = Significant model, compared via ANOVA to a null model.

Also, four ubiquitous species with notable appetence for the olive moth larvae were detected to be key predators for controlling the olive moth (see Fig. 3 and Table S1 in Appendix 2). These species were: *Tapinoma nigerrimum, T. erraticum, Aphaenogaster senilis* and *Camponotus sylvaticus*. Most ant species with a high appetence for olive moth were, however, relatively rare as to play an important role in the community predation function and pest control service (cluster of species in the upper left quadrant in Fig. 3).

**Figure 3:**
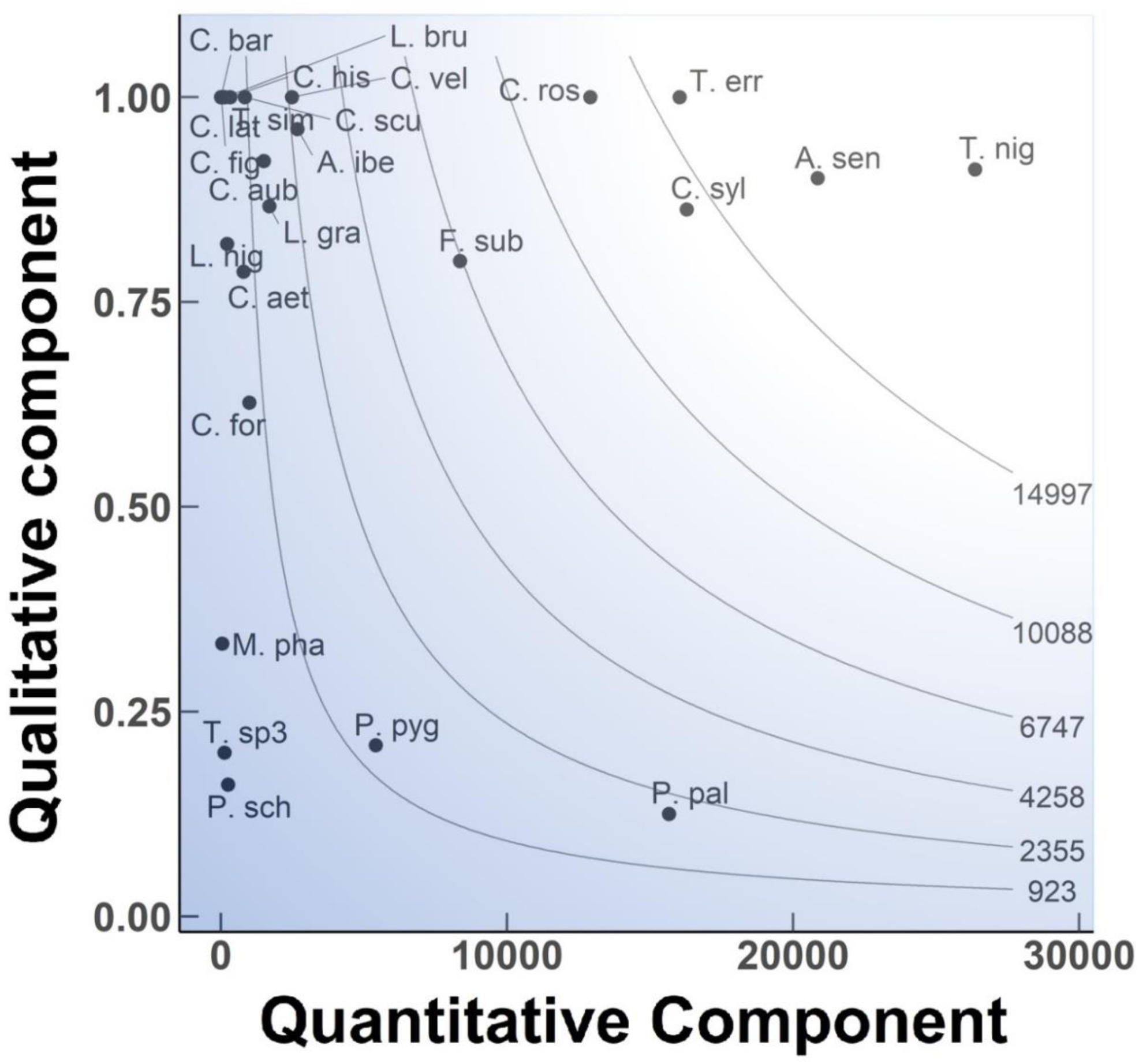
The predation effectiveness landscape of different ant species against the olive moth in Andalusian olive groves. Here, the quantitative component refers to absolute abundances. Species code correspondence to species is shown in Table S1, supporting information. Isoclines represent all possible combinations of the quantitative and qualitative components that render the same total predation effectiveness.

## Discussion

Shupp et al. (2010) introduced the dual concept of quantitative and qualitative component as two independent forces driving variations in the effectiveness of mutualistic ecosystem functions. Here, we apply and extend these concepts to enable a new framework for the quantification of the predation function, with important opportunities for the assessment of the pest control ecosystem service.

### Opportunities & Limitations

The approach shown in this study offers a new framework for scientists and practitioners to assess and compare the predation function across environmental gradients and predator groups. For instance, as our case study reveals, the predation pressure provided by ants against the olive moth increases with landscape complexity.

This approach provides an opportunity to dissect how the quantitative and qualitative components of particular species and/or communities vary through space and time, and how they respond independently (or not) to environmental gradients. This has major implications for advancing the study of antagonistic relationships, and community ecology. Since both components are not inherently correlated, divergent patterns might arise, showing what conditions maximize the predation function and proposing new questions (e.g. How? Why?). It also offers new ideas with important applications in agroecology, where maximizing the pest control ecosystem service is one of the most important challenges (Tscharntke et al., 2016). Within this framework, pest control solutions might target the optimization of the quantitative component or the qualitative component. Moreover, as exemplified in our case study, it is possible to identify key pest control agents among multiple species, and evaluate what species could be targeted to increase the predation function.

Remarkably, this is a directed, highly resolved and specific method to measure the predation function (i.e. predicts responses from and against specific species, under certain conditions). An intensive experimental or observational work is required. As a consequence, measuring species to species qualitative component in multispecies communities might be sometimes impracticable, hindering the use of this framework for wide objectives (e.g. predation pressure of birds against arthropods). The advent and generalization of the use of molecular techniques, specifically barcoding and metabarcoding, enables the identification of preys in feces, gut contents, etc., of predators, allowing the relatively rapid evaluation of the frequency of prey/pest consumption for many predators (Alberdi et al., 2020; Berry et al., 2017). Since our index may easily incorporate this kind of information for the estimation of the qualitative component of the depredation function, some of these limitations may be overcome in the near future.

It is important to note that qualitative measures are species-specific and do not account for qualitative variation between sites. For instance, the behavior or preferences of a species might vary depending on the surrounding conditions. Therefore, the qualitative factor for that species can be assessed under different conditions, or, assumed to be similar across similar sites. It is expected that predation preferences change in very different environments, but remain relatively constant in similar conditions. Scientists should decide on a case-by-case basis whether the obtained specific responses can be extrapolated to other sites or seasons. Also, there is no need to calculate specific responses for all the predator community, but a representative number of the total abundance is highly encouraged and focus should be made on a small subset of representative abundant species.

Provided similar methods for the estimation of the qualitative component, this is a standardized comparable measure, which can help transversal evaluations across systems, time and groups of organisms.

As a downside, this is an additive measure that does not account for the incorporation of intraguild predation or indirect effects, such as competitive exclusion or niche partitioning. However, these processes will be reflected on the index if they have an impact on the abundance of predators.

### Functional traits variation and the predation effectiveness framework

Functional traits can determine interactions between species and provide clues about the ecological processes underlying these relations (e.g. beak shape and feeding habits in birds) (Navalón, Bright, Marugán-Lobón, & Rayfield, 2019). The index and the components here explained can be related to functional traits at the species or community level (functional diversity or divergence) to develop mechanistic models aiming to understand variations in the predation function. For instance, the quantitative component might be related to hunting (solitary vs. group foraging) or habitat (open/forest habitats) preferences, and the qualitative component might be influenced by size (e.g. in spiders) or speed (e.g. in mammal or avian carnivores). Therefore, it is possible to define a trait-based mechanistic model of the qualitative component of the predation function, leading to accurate predictions of the predation response at the species level. Scaling up to the community level, we can subsequently calculate the community-level weighted trait and/or the trait functional divergence and use them as predictors of the community predation function and its ecological and evolutionary correlates. Some examples of such approach exist in mutualistic interactions. For example, Garrido et al. (2002) related the diaspore traits of an ant-dispersed plant and ant-wide community traits across localities, to predict seed dispersal effectiveness.

### Concluding remarks

In conclusion, understanding ecosystem functions and the derived ecosystem services provided by natural communities is key to advance our ecological knowledge and achieve sustainable agroecosystems. Here, we proposed a framework to measure the predation function accounting for the quantitative and qualitative components of interactions, which provides numerous and wide opportunities.

## Declarations of interest

none

---

**Box 1:**
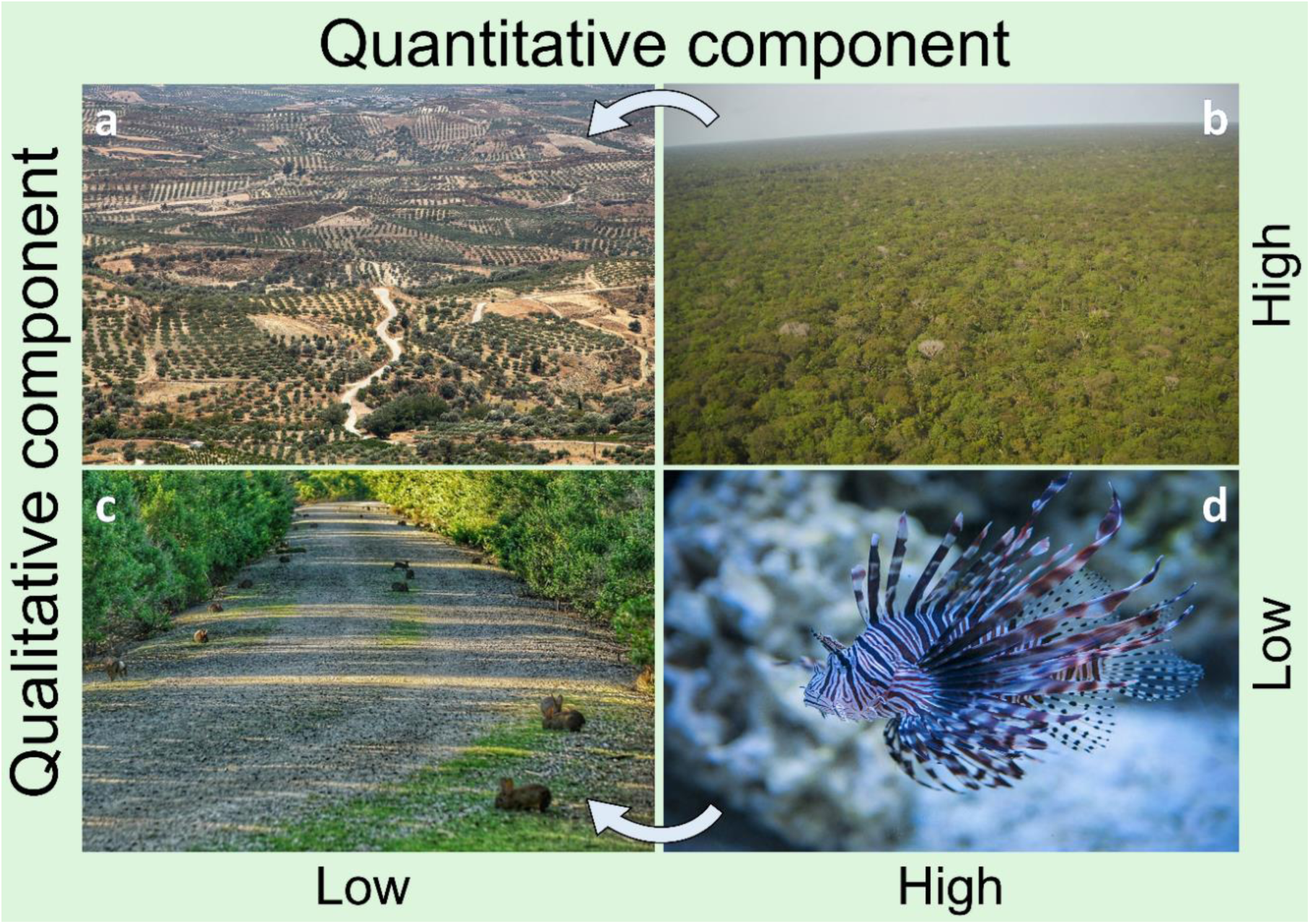
Scenarios of variation in total predation effect due to changes in the two components.

a. **High appetence for the prey (or high predation success) and low predator abundance:** This scenario is likely to be found in systems where predator abundance is constrained by biotic (e.g. competition, intraguild predation) or abiotic (limited habitat) factors. Also, in systems with highly specialized enemies or appealing prey species. The photo represents an intensive agricultural landscape with reduced remnants of natural habitat for forest insectivorous birds.
b. **High appetence for the prey (or high predation success) and high predator abundance:** This scenario maximizes the predation function, and density dependent dynamics can regulate predator and prey abundances via feedback processes. Predator-prey communities might be close to equilibrium under this scenario. The photo shows a mature stable rainforest in Brazil. Note that disturbances (arrow) might lead to *scenario a.*
c. **Low appetence for the prey (or low predation success) and low predator abundance:** This scenario minimizes the predation function, potentially leading to ecosystem function disruption. This scenario characterizes ecosystems where a non-native species has been introduced, and relatively scarce species can predate on the “new” prey. We find an extreme example in Australia with the introduction of rabbits. This introduced mammal has very few and scarce predators (e.g. dingoes or birds of prey) in Australia, that are not used to prey on rabbits, leading to the disruption of an effective predation function and the massive proliferation of rabbits.
d. **Low appetence for the prey (or low predation success) and high predator abundance:** This scenario characterizes systems where preys have evolved successful evasive mechanisms to avoid predation. Potential predators are abundant but prefer to prey on other species. For instance, the lionfish (*Ptetoris* spp.) is a venomous marine fish, native to the Indo-Pacific. Despite being little appealing to predators, there are abundant potential predators that occasionally prey on them. The lionfish is relatively scarce in their native range, but two of their species were introduced and are invasive species in the western tropical Atlantic, where they have fewer potential predators, leading to scenario c (see arrow to *scenario c*).

## Notes

### Competing Interest Statement

The authors have declared no competing interest.

