## Supporting Information for "Assessing the predation function via quantitative and qualitative interaction components"

**Title:**

[1] Dept. Biología Animal, Biología Vegetal y Ecología, Universidad de Jaén. E-23071 Jaén, Spain.

[2] Instituto Interuniversitario del Sistema Tierra de Andalucía, Universidad de Jaén, E-23071 Jaén, Spain.

Declarations of interest: none

Assessing the predation function via quantitative and qualitative interaction components

Carlos Martínez-Núñez & Pedro J. Rey

### APPENDIX 1: Methods used to calculate landscape metrics

To study how the Ant Predation Function (APF) varied with landscape complexity, we calculated six metrics of landscape complexity. Three compositional indices: PR (Patch Richness), PD (Patch Diversity), SNH (Semi-natural Habitat), and three configurational indices: LPS (Largest Patch Size), ED (Edge Density) and NND (Nearest Neighbor Distance). We used the most recent cartography of land use of the study region (www.siose.es) which was processed using Quantum GIS. Data was subsequently transferred to FRAGSTATS software ((McGarigal et al., 2012) to calculate these landscape metrics at the scale of 1 km radius around the centroid of each farm.

We ran twelve mixed model, using ant predation function as response variable, and accounting for linear and quadratic terms of the six quantitative landscape measures (one separate model for linear and other for quadratic relationships of each landscape metrics). For these models, farm ID entered as a random factor, and the ant predation function was log-transformed to meet model assumptions. Each model was compared to a null model (with no fixed factor) via ANOVA and AIC.

### APPENDIX 2

Table S1: Ant species predation function and contribution to the total predation function on the 40 olive farms, sorted from highest to lowest.

| **Species** | **Code** | **Specific response** | **Abundance** | **Total specific effect** |
|---|---|---|---|---|
| *Tapinoma nigerimum* | T. nig | 0.912 | 26355 | 24040 |
| *Aphaenogaster senilis* | A. sen | 0.901 | 20851 | 18795 |
| *Tapinoma erraticum* | T. err | 1.000 | 16033 | 16033 |
| *Camponotus sylvaticus* | C. syl | 0.863 | 16282 | 14052 |
| *Cataglyphis rosenhaueri* | C. ros | 1.000 | 12911 | 12911 |
| *Formica subrufa* | F. sub | 0.800 | 8351 | 6681 |
| *Aphaenogaster iberica* | A. ibe | 0.961 | 2672 | 2567 |
| *Cataglyphis velox* | C. vel | 1.000 | 2480 | 2480 |
| *Pheidole pallidula* | P. pal | 0.125 | 15655 | 1957 |
| *Lasius grandis* | L. gra | 0.867 | 1687 | 1462 |
| *Crematogaster auberti* | C. aub | 0.922 | 1495 | 1378 |
| *Plagiolepis pygmaea* | P. pyg | 0.209 | 5416 | 1134 |
| *Crematogaster scutellaris* | C. scu | 1.000 | 832 | 832 |
| *Camponotus foreli* | C. for | 0.627 | 989 | 620 |
| *Camponotus aethiops* | C. aet | 0.787 | 786 | 618 |
| *Cataglyphis hispanica* | C. his | 1.000 | 329 | 329 |
| *Lasius niger* | L. nig | 0.821 | 215 | 177 |
| *Lasius brunneus* | L. bru | 1.000 | 143 | 143 |
| *Tapinoma simrothi* | T. sim | 1.000 | 136 | 136 |
| *Camponotus lateralis* | C. lat | 1.000 | 67 | 67 |
| *Plagiolepis schmitzii* | P. sch | 0.161 | 248 | 40 |
| *Tetramorium* sp3 | T. sp3 | 0.200 | 127 | 25 |
| *Monomorium pharaonis* | M. pha | 0.333 | 50 | 17 |
| *Camponotus figaro* | C. fig | 1.000 | 16 | 16 |
| *Camponotus barbaricus* | C. bar | 1.000 | 4 | 4 |
